# Scaling cross-tissue single-cell annotation models

**DOI:** 10.1101/2023.10.07.561331

**Authors:** Felix Fischer, David S. Fischer, Evan Biederstedt, Alexandra-Chloé Villani, Fabian J. Theis

## Abstract

Identifying cellular identities (both novel and well-studied) is one of the key use cases in single-cell transcriptomics. While supervised machine learning has been leveraged to automate cell annotation predictions for some time, there has been relatively little progress both in scaling neural networks to large data sets and in constructing models that generalize well across diverse tissues and biological contexts up to whole organisms. Here, we propose scTab, an automated, feature-attention-based cell type prediction model specific to tabular data, and train it using a novel data augmentation scheme across a large corpus of single-cell RNA-seq observations (22.2 million human cells in total). In addition, scTab leverages deep ensembles for uncertainty quantification. Moreover, we account for ontological relationships between labels in the model evaluation to accommodate for differences in annotation granularity across datasets. On this large-scale corpus, we show that cross-tissue annotation requires nonlinear models and that the performance of scTab scales in terms of training dataset size as well as model size - demonstrating the advantage of scTab over current state-of-the-art linear models in this context. Additionally, we show that the proposed data augmentation schema improves model generalization. In summary, we introduce a de novo cell type prediction model for single-cell RNA-seq data that can be trained across a large-scale collection of curated datasets from a diverse selection of human tissues and demonstrate the benefits of using deep learning methods in this paradigm. Our codebase, training data, and model checkpoints are publicly available at https://github.com/theislab/scTab to further enable rigorous benchmarks of foundation models for single-cell RNA-seq data.

## Introduction

Cell type annotation is a core step in the analysis of single-cell RNA-seq (scRNA-seq) data. Researchers typically examine prominent gene expression markers denoting a cell’s identity and function, and assign a label based on a nomenclature that summarizes previously described cell types and states. While this task has been addressed in numerous analyses^1–4^ and automated cell type prediction models^5–8^, rigorously annotating new datasets remains a manual and time-consuming process. Moreover, given the confounding presence of technical batch effects and the inherent differences in quality across cells within datasets, the process of generating cell annotations remains unstandardized. These problems became especially pronounced in building comprehensive cell atlases in the Human Cell Atlas (HCA)^9^, wherein unannotated datasets remain a bottleneck. Indeed, recent atlas-building efforts acutely highlight the challenges posed by both the lack of consensus in cell type annotations across datasets and how time-intensive it remains to standardize them^10,11^. A general model for cell type annotation predictions — that is, a model trained on a large and diverse data corpus consisting of all human tissues in diverse — would assist with the atlas-building efforts of the HCA in several crucial ways: To begin with, such a model would lower the barrier of manually annotating datasets for researchers, offering suggestions with a standardized set of nomenclature. Predictions with a uniform set of vocabulary will naturally push the community to adopt consistent terms when referring to cell types. Moreover, such model suggestions will serve as “hints”, a baseline for researchers to modify and refine based on their own knowledge and expertise. Finally, such a process would allow scientists to annotate datasets at the scale required by the HCA and related initiatives.

Providing cell labels for unannotated datasets can be posed as a machine-learning classification task. However, cross-tissue classifiers trained on large-scale data collections that annotate cells from heterogeneous sources, irrespective of tissue of origin and assay type, are surprisingly slow to emerge. Models often address only specific scenarios and do not focus on strong generalization capabilities beyond the datasets they are trained on^12^, partly due to fragmented data collection efforts. Recent large data curation efforts streamline the training of generalist models because they expose cell type labels and other metadata in structured vocabularies of ontologies and consistent feature spaces^13,14^. In particular, CELLxGENE hosts a first draft of a curated data collection that allows for models to be trained across a significantly larger range of datasets than was possible before^15^. Nevertheless, a dominant paradigm of the cell type annotation task has been annotation transfer^16^, specifically the approach of projecting entire samples of cells on an annotated reference atlas to transfer cell type labels^1,17,18^. Annotation transfer is often used in organ- or lineage-specific scenarios, yet it is critically limited by the quality and similarity of the annotated reference. In contrast to this paradigm of query-to-reference mapping, we focus on the problem of general cell type annotation which entails training models across a large-scale data corpus to predict cell type labels solely based on gene expression. It’s also worth noting that the predictions of such a model would be community-driven; researchers would not be required to rely upon annotation transfer-based methods. Such approaches inevitably suffer from model overtraining upon a single reference (often with lab-specific cell labels) and strongly encourage researchers to choose a context-specific reference close enough to their study of interest as a basis for trustworthy cell annotations.

Several aspects of the general cell type annotation problem remain ambiguous: Firstly, initial attempts to increase model complexity in cell type annotation to improve classification performance have failed to improve over linear baseline models^6,19,20^. Consequently, the question pertains to whether large-scale, non-linear models learn cell state representations that are more useful for this classification task than linear, well-tuned baseline models that are trained on large-scale datasets as well. Furthermore, recent efforts have started to use large-scale data corpora with tens of millions of scRNA-seq profiles to train deep learning models^21–23^. However, those efforts on foundation models benchmark cell representations on diverse tasks, so far without context on deep models specifically designed for cell type annotation. Indeed, the cell annotation tasks considered in these efforts are either only fine-tuned to specific cell type classification problems with only a few cell types^21,22^ or do not directly predict cell type labels but rely on finding similar cells in an annotated reference^23^. Moreover, these initial attempts at building foundation models that cover a large data corpus largely treat the cell type labels as mutually exclusive and ignore label relations that were previously exploited in organ-centric classification tasks^14,24^, thus questioning if the resulting benchmarking metrics are faithful evaluations of the performance on these heterogeneous datasets. Lastly, it remains unclear how cross-tissue cell type classifiers compare their respective organ-specific counterparts. Here, we address these challenges by assembling a benchmark dataset for cross-tissue cell type classification and carefully analyzing a cross-tissue cell type classifier optimized for cell type annotation on tabular scRNA-seq data: scTab. scTab uses observation-wise feature attention to reduce the number of input features for each observation. This helps the model to be more robust to overfitting to poorly generalizable features in the training data, which is often an issue for tabular data as there is no prior knowledge about the underlying structure of the data - unlike for e.g. images or text^25^.

We leverage well-defined benchmark metrics^5,7^ for cell type classification to understand the performance of deep learning models trained on large scRNA-seq data corpora, focusing on the scaling behavior of such models with respect to the training data size as well as the model size^26^. We find that analogous to the work in computer vision, cell type classification from scRNA-seq data substantially benefits from large-scale training of deep-learning-based models^27^, and model generalizability can be improved by artificially increasing the training data size through data augmentation^28^. In addition, we find that by scaling cell type classification to large-scale datasets, deep-learning models outperform their linear counterparts, in contrast to what was reported before^6^. Yet, well-defined baseline models are still relatively powerful, thus suggesting caution in the design of benchmarking experiments in foundation models. In summary, our detailed analysis demonstrates the strengths of deep learning-based approaches over their linear counterparts in large-scale, cross-tissue cell type classification and shows that classification performance scales both with respect to training data and model size.

## Results

### A dataset and evaluation metric to study the scaling behavior of cross-tissue cell type classification models

We set out to build a dataset on which a cross-tissue cell type classification model could be trained and evaluated. In the existing literature, we identified three approaches to creating such a dataset. The first approach involves assembling study- or organ-specific datasets and homogenizing cell type labels in order to obtain a mutually exclusive set of labels or a tree with levels of mutually exclusive labels^5–7^. The second approach centers around assembling organ-specific datasets and mitigating annotation granularity differences by using the Cell Ontology^29,30^ to establish a directed acyclic graph between the observed labels^14^. The third approach entails the collection of an organism-wide data corpus with ontology-constrained labels treated as mutually exclusive, thus ignoring the hierarchical structure of labels in the ontology^21,22^. This is problematic since the hierarchical dependency between labels is a key structure of these datasets, e.g. a “CD4-positive, alpha-beta T cell” is also a “T cell” and a “lymphocyte”, and penalizing a model that predicts “CD4-positive, alpha-beta T cell” for a cell that is labeled as a “T cell” results in an evaluation that is biased towards the model being able to mimic the annotation granularity of the data instead of its ability to distinguish cell types. Here, we leveraged the cell type relations given by the Cell Ontology^29^ across a data corpus of all human tissues, using a release of the cell census by CELLxGENE as a root dataset (Methods). This data corpus reflects a large number (164) of cell types in the human body, therefore, we refer to the resulting label prediction problem as “general cell type classification”.

To benchmark a cell type prediction task that represents the recent literature closely^7,20^, we considered models with a softmax-constrained output across all observed cell type labels. To account for differences in annotation granularity across datasets, we adjusted predictions based on the Cell Ontology to not incur penalties if a model predicts a more fine-grained label than the original author annotation (Methods). We modified the original release of the cell census in preparation for this task (Methods). First, it is important to realize that public data corpora are not necessarily deduplicated. Often, cells are present in a primary dataset that originates from an original study and are also contained in secondary datasets (metastudies), such as atlas datasets. We only considered instances of cells in primary datasets to avoid data leakage through duplicate cells. Second, we removed cells with broad cell type labels, using a heuristic that removes each cell type with less than seven parent nodes in the Cell Ontology. Third, we restricted to cells measured with the most common group of 10X Genomics technology-related assays, to reduce the strength of confounding sources of variation in the dataset. Fourth, we removed cells from rare cell types with less than 5,000 instances or those present in less than 30 donors to accurately assess how the trained classifiers generalize to unseen donors (Methods). The resulting dataset contains 22.2 million cells, with 5,052 donors and 164 cell type labels (Fig. 1a). We defined test holdouts based on donor annotation, which we see as a sensible compromise between an entirely random split and a split based on holdout studies. The donor-wise split improves the coverage of labels in both training and test sets compared to a split based on studies, and reduces leakage of similar observations between training and test data compared to a random split of cells, thus creating an evaluation set that is better suitable to assess the generalization capabilities of a classifier. As a benchmarking metric, we chose the macro-averaged F1-score (macro F1-score) (Methods) to account for class imbalances and to give each cell type an equal weight in the overall score.

**Figure 1:**
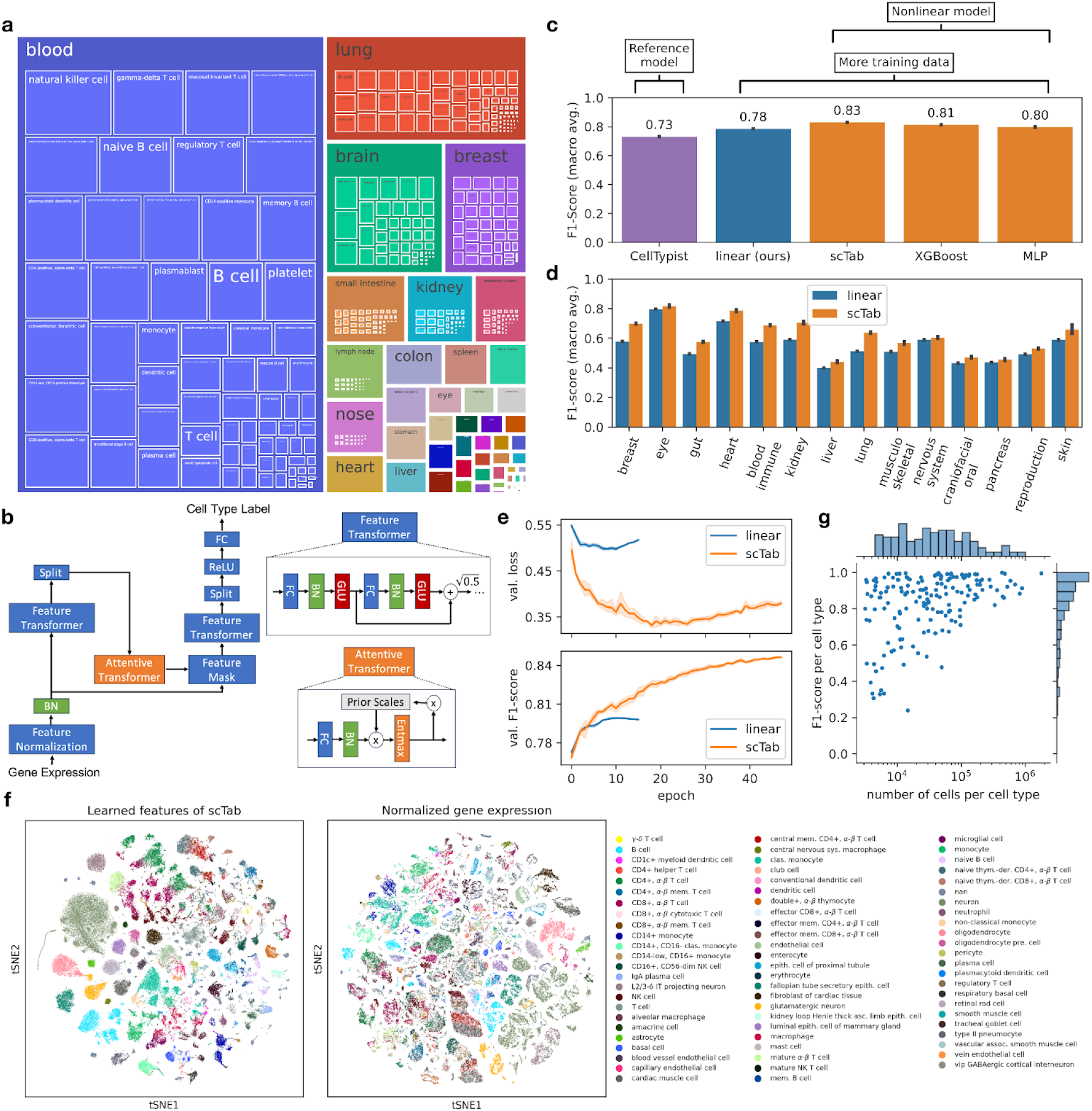
scTab enables organism-wide, scalable, and robust cell type classification on single-cell RNA-seq data. **(a)** Treemap plot showing the composition of the assembled dataset across cell types and tissues. Each color corresponds to one tissue with the size of the box giving the number of donors for that tissue and the inner boxes the number of donors for each cell type. The 22.2 million cells from the assembled dataset span 5,052 unique donors, 249 datasets, 52 disease states, 164 unique cell types, and 56 different tissues. A full list of the cell type label, tissue, and number of donors and cells per combination of label and tissue is given in Supp. Table 5. **(b)** Overview of the scTab architecture used in this paper (Methods). After normalization of the input features (gene counts are normalized to 10,000 counts per cell and then lop1p transformed), it encodes data via a feature transformer and selects relevant input features through feature attention via an attention transformer block. (FC: fully connected layer, BN: batch norm layer, GLU: gated linear unit nonlinearity, ReLU: rectified linear unit) **(c)** Comparison of classification performance (measured by macro F1-score) of linear reference models (CellTypist (retrained on cross-organ data and subsampled to 1.5 million cells), Linear) and nonlinear models (scTab, XGBoost, MLP (multi-layer perceptron)). **(d)** Classification performance (measured by macro F1-score) grouped by organ system of scTab and our linear reference model. **(e)** Cross-entropy loss and macro F1-score on the validation set plotted after each epoch for scTab and our linear reference model. The performance difference between the two models is stronger for the cross-entropy loss than for the macro F1-score. **(f)** tSNE plots of raw features (input to linear classifier) and the learned features of scTab with the top 70 most frequent cell types superimposed. Plots show the holdout test data. **(g)** F1-score per cell type plotted against the number of unique cells observed per cell type for scTab. The histogram on the y-axis shows the distribution of F1-scores and the histogram on the x-axis shows the distribution of unique cells per cell type (log scale).

### A feature-attention-based, scalable, deep-learning model for cross-tissue cell-type classification

Studying the scaling behavior of deep-learning-based models necessitates a scalable model implementation that can be trained on bigger-than-memory datasets. In addition, we ask if recent extensions beyond classical multi-layer perceptrons (MLP) improve prediction as they have in other fields. Since gene expression profiles are not ordered, we decided against sequence-based models such as transformers^21,22^ and instead selected a recent architecture specifically proposed for tabular data^31^. Here, we introduce scTab (Fig. 1b), which is a scalable implementation of the TabNet architecture^31^, which we adapted to the single-cell use case: scTab is specifically designed for the tabular structure of scRNA-seq data through the use of feature attention, which enables the network to focus its model capacity on more reliable input features. After normalization, it encodes data via a feature transformer and selects relevant input features through feature attention via an attention transformer block (Methods). We modified the original TabNet implementation in a few crucial ways: scTab’s input data assumption is adapted to the single-cell setting, in particular, the input gene expression is size factor normalized to 10,000 counts per cell and log1p transformed. This common normalization for scRNA-seq data^7,23^ cannot be replicated by the simple batch normalization layer used in the original TabNet architecture. We additionally modified the original TabNet architecture to improve computational efficiency, namely by reducing the number of feature and attention blocks (which we found unnecessary after profiling), and training dynamics for faster convergence (Methods). For better model generalizability, we further added a data augmentation step as described later below. Finally, scTab quantifies prediction uncertainty using empiric uncertainty probabilities based on deep ensembles^32^ (Methods).

### Cross-tissue cell type classification requires nonlinear models

To showcase the performance of state-of-the-art models according to recent benchmarks^5,20^, we first retrained a CellTypist model^7^ (Methods) to a random subsample of our training corpus. The current CellTypist implementation necessitated the full training data to be subsampled for model training as on the one hand it requires all the training data to be loaded into memory and on the other hand it lacks GPU acceleration. Here, we subsampled to 1.5 million cells. This re-trained reference model achieved a macro F1-score of 0.7304±0.0015 (± is indicating the standard deviation) (Fig. 1c). Given this performance of the reference CellTypist model, we investigated if performance could be increased by scaling logistic regression-based models to take advantage of the full training data size. We implemented a logistic regression-based model not subject to these limitations (Methods) and trained this model with a cross-entropy loss. This model significantly outperformed the CellTypist reference model and achieved a macro F1-score of 0.7848±0.0001 (Fig. 1c), showing the potential of scaling existing linear models to take advantage of larger datasets. Having optimized the linear reference model, we benchmarked three nonlinear models against this baseline: our scTab model, an MLP previously proposed for this task^6,14^ - but found to not outperform linear models within single tissues - and an XGBoost model that reported robust performances on classification tasks for tabular data^25^: The nonlinear models significantly outperformed the linear model (0.8295±0.0007 macro F1-score for scTab (fitted with data augmentation), 0.8127±0.0005 for XGBoost, 0.7971±0.0012 for MLP (fitted with data augmentation)) (Fig. 1c, Supp. Table 1), demonstrating that cross-tissue cell type classification is complex enough to benefit from nonlinear models. Moreover, scTab outperformed the linear model on all organ systems when these were considered separately (Fig. 1d). Further to this, when looking at uncertainty scores calculated based on deep ensembles^32^ (Methods), one can see that incorrect predictions on the holdout test data are also associated with a higher model uncertainty (Supp. Fig. 5). Besides the performance advantage over linear models, scTab also showcases different training dynamics: On the one hand, the difference to the linear model is more pronounced when looking at the loss, and on the other hand, scTab is trained for more epochs (Fig. 1e). When qualitatively inspecting the representations learned by scTab in a tSNE plot, we found that cell types show consistent separation and that the latent space was more structured compared to the raw feature space (Fig. 1f). In summary, we show that leveraging larger and more diverse training data sets coupled with deep learning-based models dramatically improves classification performance for cross-tissue cell type classifiers (Fig. 1c). Furthermore, one can see that classes on which errors were made tended to be those that were represented by few observations (Fig. 1g). This further motivates that classification performance can indeed be improved by adding more training examples specifically for cell types which are hard to classify.

### Cross-tissue cell type classification scales with dataset and model size

A key driver behind the success of deep-learning-based models in computer vision or natural language processing is their ability to take advantage of larger datasets. This scaling behavior was a driving factor of recent advances in computer vision and natural language processing and led to the study of how model performances scale with model size and training examples^26,27^. Having established that the cross-tissue cell type classification problem satisfies this premise, we next set out to study its scaling behavior. In contrast to images, the heterogeneity of a scRNA-seq corpus is not trivially measured by the number of observations (cells). Some datasets contain densely sampled cell states, in which new samples would simply replicate what is already captured, whereas other samples are relatively sparsely sampled (Fig. 2a). To account for this complexity in the study of data scaling behavior, we compared the test performance of models trained on subsets of the full corpus, either subsampled randomly by cells as a control, or subsampled by donors to tie the subset size closer to the relative complexity captured by this dataset (Methods). Indeed, we observed a strong scaling of the loss and macro F1-score with respect to the dataset size for donor sub-sampling, but a much weaker scaling for cell-subsampling (Fig. 2b) indicating that scaling with respect to the training dataset size is mostly driven by batch diversity rather than the number of cells. This scaling also held for all organ systems when inspected individually, with a minimum difference in macro F1-score of 0.0454±0.0083 between a dataset of 2.1 million cells and the full training dataset of 15.2 million cells, and a median difference of 0.1219±0.0212 (Fig. 2c). A potential reason for the difference in classification performance between different organ systems lies in the number of observed cell types per organ system - the F1-scores have a correlation of -0.55 with the number of observed cell types per organ system. The observed scaling with data size was also specific to scTab and was not exhibited by the linear model (Fig. 2d), whose learning curve flattens out earlier, resulting in an improvement of the macro F1-score by scTab over our linear reference model of 0.0447±0.0008 when using the full training data. Moreover, we compared the performance of lung-specific cell type classification of models trained only on lung data against their cross-organ counterparts. The deep-learning-based scTab model shows more robust performances compared to its linear counterpart (Fig. 2e). The lung-specific performance of scTab only drops from a macro F1-score of 0.7220±0.0078 to 0.7062±0.0122, whereas the performance of the linear model drops from 0.7146±0.0040 to 0.5291±0.0041 (Supp. Table 2), suggesting that cross-organ cell type classification benefits from using non-linear models. This also indicates that if the use case is purely organ-specific, adding this information as an additional covariate might help the classifier achieve better classification performance; hence a simple extension of scTab would be to add the tissue as an additional input covariate.

**Figure 2:**
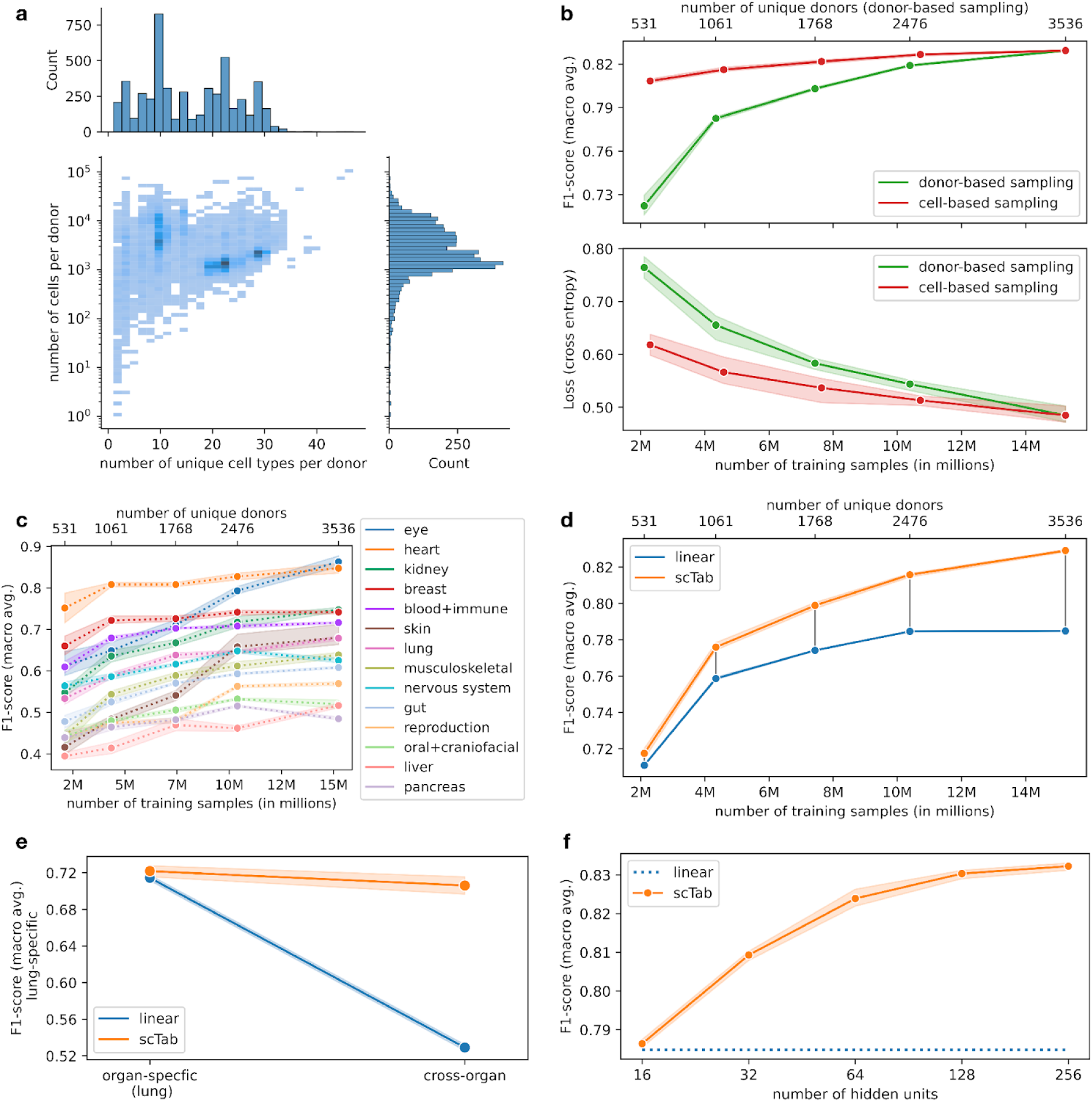
Non-trivial scaling behavior of scTab in cross-tissue cell type prediction. **(a)** Distribution of donors with respect to the number of unique cell types (x-axis) and with respect to the number of cells (y-axis). The histogram on the y-axis shows the distribution of donors with respect to the number of cells (log scale). The histogram on the x-axis indicates the distribution of donors with respect to the number of unique cell types. **(b)** Scaling behavior of scTab with respect to the size of the training data for two simulated scenarios in terms of macro F1-score and cross-entropy loss: i) cell-based subsampling which corresponds to increasing the number of sequenced cells while keeping the observed biological diversity constant ii) donor-based subsampling which corresponds to increasing the observed biological diversity and which also reflects the real world more closely as new datasets will mostly contain new unobserved donors. All cell types from the test set were observed during model training for all subsampled datasets. **(c)** Scaling of the cross-organ model from Fig. 2b with respect to training data size grouped by organ system (subsampling is done based on donor-based subsampling). **(d)** Scaling behavior of scTab versus our linear reference model with respect to the training data size. One can see that the difference between the two models increases with increasing training data size. **(e)** Effect of training only on lung-specific data versus training on all cross-organ data on lung-specific classification performance (evaluated on test data subset only to lung data) for scTab and our linear reference model. **(f)** Scaling behavior with respect to model size. The number of hidden units refers to the size of the fully connected layers (FC) in the architecture (Fig. 1b, Methods).

Focussing on assessing model capacity at increased data size, we performed a second scaling experiment in which we kept the full dataset but compared scTab implementations with different numbers of parameters (Methods). We found a significant improvement in performance between the smallest model with 1.7 million parameters (0.7864±0.0010 macro F1-score) and the largest model with 16.2 million parameters (0.8323±0.0010 macro F1-score) (Fig. 2f). Overall, the above-mentioned differences in scaling behavior to a baseline linear model show that a nonlinear model is better able to take advantage of larger and more diverse data sets and that it is able to model complex non-linear relationships.

### Data augmentation improves classifier generalizability

Due to their high model capacity, deep learning models are known to easily overfit the training data and can even fit random labels^33^. One well-established technique to reduce overfitting and thus improve model generalizability is to artificially increase training data size by applying semantically-preserving transformations. For images, these transformations include rotating or cropping input images during training^28^. Data augmentation serves as a regularization technique that yields models with better generalization capabilities and less impacted by dataset-shift phenomena. So far, data augmentation has not been consistently applied in single-cell genomics, due to the limited capacity of most scRNA-seq models, and due to the lack of sensible augmentation strategies. Here, we propose a novel augmentation strategy for scRNA-seq data and evaluate it for cell type prediction with our scTab model. Notably, our data augmentation strategy is not only limited to scTab but can be used in combination with other models as well. The motivation for the proposed data augmentation is to simulate the gene expression vector of a target cell if it were observed in a different donor. To do so, we precompute augmentation vectors based on the training data that can be added to the original gene expression vectors during model training (Fig. 3b). The data augmentation vectors are the average difference computed in the full gene space between cells of the same cell type observed in two different donors. Thus, by adding those augmentation vectors to the gene expression vector of the original cell, one can simulate the gene expression vector of a target cell in a different donor, extending the training data domain in these incompletely observed donors (Fig. 3a). Before evaluating the effect of this augmentation on model fits, we established that it did not severely disrupt the training data structure. Boundaries between cell types are blurred in a tSNE of the augmented data. Still, cell type identity as a main source of variation in the data is preserved (Fig. 3c) as quantified by a similar variance decomposition in terms of cell type and donor labels (𝑅^2^=0.189 for the raw data, 𝑅^2^=0.164 for the augmented data, Supp. Table 7, Methods). We found this augmentation strategy to regularize models, training loss increased upon using augmentation, and the macro F1-score on the training data decreased. Model generalization was improved on the validation set as measured by reduced loss and increased macro F1-score (Fig. 3d). When looking at the holdout test set, the proposed data augmentation significantly reduces the loss from 0.797±0.05 to 0.659±0.04 (p-value: 0.0039) and significantly increases the macro F1-score from 0.7755±0.0020 to 0.7841±0.0030 (p-value: 0.0016) (Supp. Table 3). These results show that sensible data augmentation techniques for scRNA-seq data can significantly improve the generalization performance of cross-tissue cell type classifiers.

**Figure 3:**
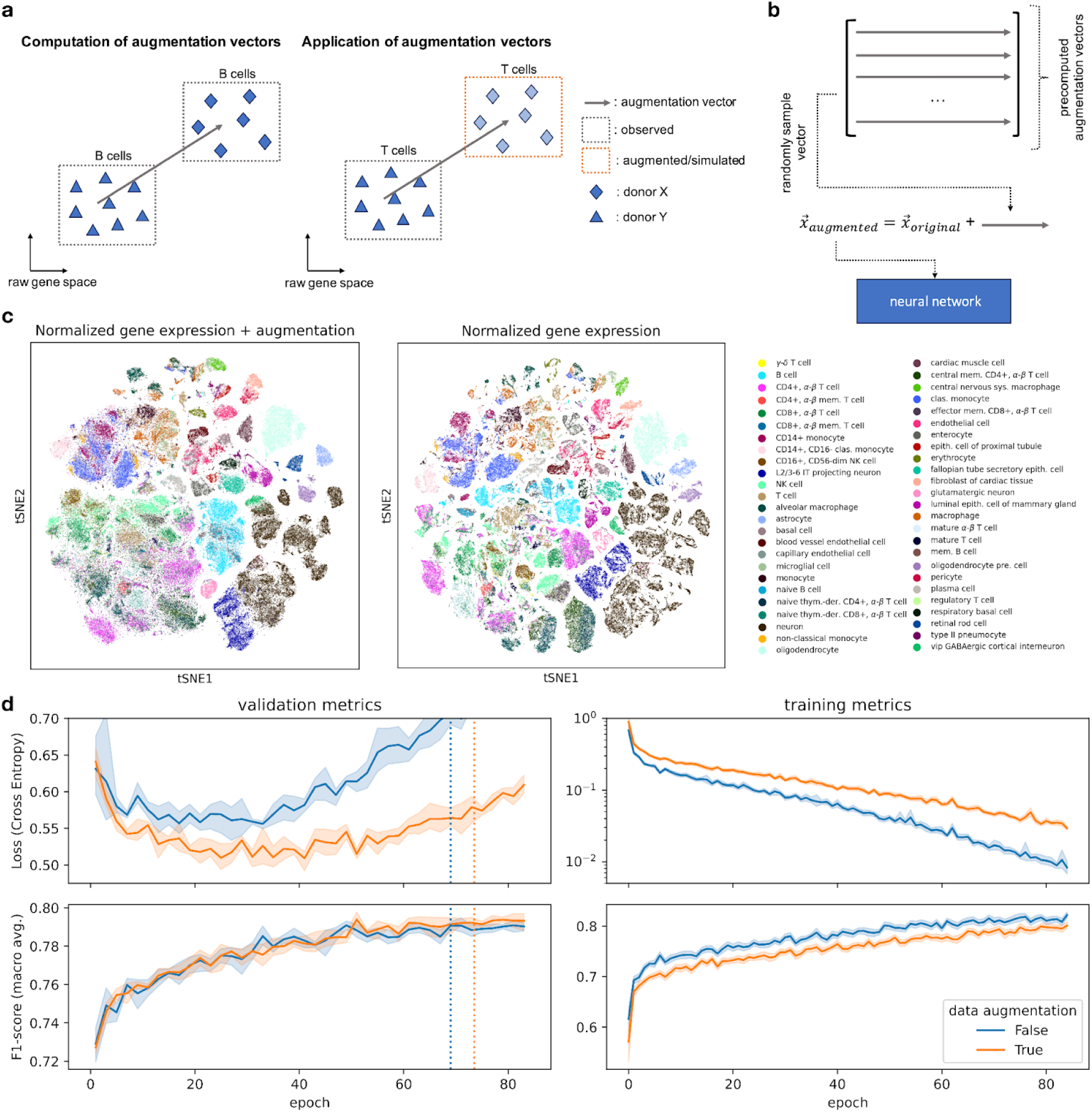
Data augmentation for scRNA-seq cell type classification improves model generalizability. **(a)** Illustration of the data augmentation procedure. The difference vector in raw gene space between the same cell type observed across two donors can be used to simulate how the gene expression of a cell type might look for a different donor and, thus, artificially increase the training data size. (**b)** For each input vector to the neural network, an augmentation vector is randomly sampled and added to the original input vector. The augmented vector is then fed into the neural network (due to simplicity the batch dimension is omitted in the sketch). **(c)** tSNE visualization of original and augmented data. One can see that the augmentation blurs out the boundaries of the cell types but that the main source of variation (cell type) is still preserved. **(d)** Effect of augmentation on training and validation loss and macro F1-score (training data was subset to 4.3 million cells (Methods)). One can observe the desired effect of data augmentation, an increase in training loss (regularizing effect), and a decrease in validation loss. The dashed vertical lines indicate how long the models with and without data augmentation are fitted on average (early stopping is done based on the macro F1-score), respectively.

### Robust benchmarks for cross-tissue cell type classification

It is common practice in machine learning to have standardized large-scale benchmark data sets such as the ImageNet^34^ subset for the “ImageNet Large Scale Visual Recognition Challenge”^35^ and the Microsoft COCO dataset^36^ in computer vision, the GLUE/SuperGLUE dataset^37,38^ and the WMT2014 English-German datase^39^ in natural language processing. These benchmark datasets enable models to be trained on bigger data corpora and allow for structured model benchmarks that usually do not require re-training of reference models. Such ready-to-use and large-scale benchmark datasets for cell type classification on single-cell transcriptomics data are not yet easily accessible. Creating such datasets for scRNA-seq data comes with two key challenges: On the one hand, such datasets need to come with a performant and easy-to-use data-loading infrastructure, that is able to scale to bigger-than-memory datasets. Otherwise, it becomes challenging for users without the proper technical background to use such datasets in their workflow. On the other hand, such datasets should be predefined, easily accessible, and come with fixed training, validation, and test splits to make results easily comparable. Now, to encourage similar practices, our processed benchmark dataset with predefined train, validation, and test splits and the accompanying data loading infrastructure are available to download (Methods). The downloadable dataset is ready to use out-of-the-box with an efficient data loader (Supp. Fig. 1, Methods). Furthermore, the dataset comes with a set of well-tuned reference models (Methods) that can be directly used for further benchmarking efforts. The need for well-tuned reference models is demonstrated by the comparison of the performance of the XGBoost and CellTypist models given default parameters and their respective performance given tuned parameters. On the benchmark data, the performance, measured by macro F1-score, could be increased from 0.5855±0.0112 to 0.8127±0.0005 for the XGBoost model and from 0.6258±0.0036 to 0.7304±0.0015 for the CellTypist model respectively (Supp. Table 4). We expect this combination of a well-defined benchmark dataset with well-tuned baseline models to facilitate the systematic study of model scaling laws, which are of importance for the establishment and evaluation of foundation models^21–23,40,41^.

## Discussion

We introduced cross-tissue cell type classification on a whole-body human data corpus of scRNA-seq data as a machine learning task that facilitates cell type annotation and that can benefit from large-scale data collections and the usage of larger, non-linear models similar to examples in computer vision^27^. Notably, even on a well-defined dataset and with optimized models for tabular data, this task is not yet perfectly solved. We demonstrated scaling of model performance with training dataset size and model size on this task, noting that batch diversity dominates the raw number of cells in this data scaling. We also found that model overfitting can be mitigated through data augmentation. Additionally, the analysis and models introduced here provide a reproducible context for future work on cross-tissue cell type classification which is a cornerstone in the context of foundation models for scRNA-seq data, for example, by providing a standardized large-scale benchmark dataset and a set of well-tuned reference models.

General cell type classification reflects the ability of models to learn cell types based on transcriptomic profiles, a key abstraction of scRNA-seq data. But, like many supervised machine learning tasks, it is limited by the annotation granularity of the training data. The CELLxGENE data corpus used here is based on the cell ontology. As the Cell Ontology is still a work in progress, not all cell types can be correctly matched to a corresponding ontology term, this is especially a problem for rare cell types. Moreover, relationships between cell types given by the Cell Ontology are still a topic of active discussion, which can affect the classification metrics discussed here. However, we would like to highlight that the strength of these current models for automated cell type annotation does not lie in correctly classifying novel cell types, but rather in context-specific suggestions which biologists can further refine. Besides, the models from our paper can be readily retrained once more and better-annotated data becomes available.

Future work may extend the concept of general cell type classification to less stringent filters of the public data corpus, for example including cells from assay technologies that are not as common as the technologies considered here, and including rarer cell types. We would like to highlight that those efforts will need to take particular care in defining more detailed evaluation metrics, as plain macro F1-scores may not properly reflect the complexity of extremely unbalanced datasets with 100s-1000s of classes. In addition, we would like to emphasize that the performance of machine learning models can be heavily influenced by the composition and quality of the training data. For example, by specifically collecting more training data for cell types or tissues a model struggles with or by being more rigorous with the training data selection through only selecting datasets with high-confidence annotations. Here, we would like to stress that predictions will become more refined, once more refined training data becomes available; and note that the growing magnitude of single-cell data is not limited to transcriptomics; single-cell researchers have increasingly utilized spatial transcriptomics, proteomics, and other multimodal assays to investigate how other features (e.g. chromatin accessibility, DNA methylation, etc.) could be used to distinguish between cell types and cell states. Building on this point a future direction of work would be to extend scTab to take advantage of different input modalities as well once data for those modalities becomes available at an equally large scale, exploiting this feature space to achieve more precise cell type predictions for cell identities. Furthermore, recent efforts to establish foundation models for scRNA-seq data used further tasks to characterize their ability to learn nontrivial representations of cells. We envision further benchmarks to individually address these specific tasks, again focussing on data and strong baseline models. In this context of cellular representation learning, further and more refined choices for data augmentation may be explored. Additionally, these augmentation schemes can then be evaluated in the context of unsupervised representation learning like for example Bootstrap Your Own Latent^42^.

Finally, it is critical to make general cell type classification models like scTab easily accessible to the broader community of biological researchers. The Cell Annotation Platform (CAP; https://celltype.info/) has been specifically designed for HCA researchers of all backgrounds to effectively work with the predictions of scTab (as well as other prediction algorithms) directly via their browsers. With the promise of high-confidence predictions of cell types and cell states using a structured vocabulary, as well as the ability to refine and edit these predictions or reannotate cells entirely, researchers will be empowered to annotate their datasets at the current scale required to construct large-scale human cell atlases.

## Methods

### Dataset preparation

The dataset used in this paper is based on the CELLxGENE^15^ census version 2023-05-15 (https://chanzuckerberg.github.io/cellxgene-census/index.html). The census version 2023-05-15 is selected as it is a long-term supported (LTS) release and will be hosted by CELLxGENE for at least 5 years. This makes the dataset creation easily reproducible for the foreseeable future. We subsetted to human datasets and used the human protein-coding genes (19,331) as a feature space.

The following criteria are used to filter the human CELLxGENE census data:

1. The census data is subset to primary data only (*is_primary_data == True*) to prevent label leakage between the train, validation, and test set.
2. Only sequencing data from 10x-based sequencing protocols is used. In terms of the CELLxGENE census, this means subsetting the *assay* metadata column to the following terms: 10x 5’ v2, 10x 3’ v3, 10x 3’ v2, 10x 5’ v1, 10x 3’ v1, 10x 3’ transcription profiling, 10x 5’ transcription profiling
3. The annotated cell type has to be a subtype of the *native cell* label based on the underlying cell type ontology
4. For each cell type, there have to be at least 5,000 unique cells. Otherwise, the whole cell type is dropped from the dataset.
5. Each cell type has to be observed across at least 30 donors to reliably quantify whether the trained classifier can generalize to new unseen donors for each cell type. With the used 70-15-15 train, validation, and test split this means that each cell type is represented with at least 4-5 donors in the validation and test set, respectively.
6. Each cell type needs to have at least seven parent nodes in the cell type ontology. This criterion is used as a heuristic to filter out general cell type labels that do not contain much information.

To be able to better assess how well the trained classifiers generalize to unseen donors or in general to better assess the generalization capabilities of the trained classifiers, the data is split into train, validation, and test sets based on donors and not based on random subsampling. Meaning, each donor is exclusively found either in the training, validation, or test set. Unlike splitting based on e.g. holdout datasets, donor-based splitting mostly preserves the proportion of cells in the training, validation, and test set compared to random subsampling. This is not the case when subsetting the available data based on e.g. datasets, which often results in a very uneven distribution of cells across the training, validation, and test sets as the datasets in the census usually range anywhere between a few thousand cells to a few million cells. Furthermore, dataset-based splitting often makes it hard to ensure that each cell type is observed across both the training data as well as the test data. In the end, the data is split such that 70% of the donors are assigned to the training set and 15% of the donors are assigned to the validation and test set respectively.

The data is size factor normalized to 10,000 counts per cell and log1p-transformed.

The selection described above results in 22,189,056 cells being selected which span 164 unique cell types, 5,052 unique donors, and 56 different tissues. Of the 22.2 million cells 15,240,192 cells are assigned to the training set, 3,500,032 are assigned to the validation set and 3,448,832 cells are assigned to the test set.

More detailed explanations and references to the code that can be used to reproduce the above data selection and splitting exactly can be found in the associated GitHub repository under *docs/data.md*.

### Subsampled datasets

We used a subsampled training dataset in the following settings:

Dataset size scaling:

● Random subsampling: 15% subsampling (2.3 million cells), 30% subsampling (4.6 million cells), 50% subsampling (7.6 million cells), 70% subsampling (10.7 million cells), 100% subsampling (15.2 million cells)
● Donor-based subsampling: Subsample to 15% of donors (531 donors / 2.1 million cells), Subsample to 30% of donors (1,061 donors / 4.3 million cells), Subsample to 50% of donors (1,768 donors / 7.4 million cells), Subsample to 70% of donors (2,476 donors / 10.4 million cells), Subsample to 100% of donors (3,536 donors / 15.2 million cells)

Data augmentation:

- Subsample to 30% of donors (1,061 donors / 4.3 million cells) In all other cases, the full training dataset is used.

All subsampling is done incrementally, e.g. the 30% subsampled dataset includes all cells/donors that are present in the 15% subsampled dataset and so forth.

### Data loading infrastructure

Training machine learning models on large-scale tabular datasets (which is the case for the scRNA-seq data used in this paper) comes with a set of unique challenges. The first challenge is that the entire dataset does not fit into the memory of a usual server commonly used for training deep learning models. Additionally, the unique nature of tabular data means that you cannot load individual observations from disk efficiently, as individual observations are rather small, and thus loading data points individually creates a lot of random reads which even modern SSDs cannot handle efficiently. Thus, a consecutive block of samples must be loaded at once and then shuffled. Fortunately, there already exist Python libraries that do exactly what is described above. The data loading infrastructure used in this paper is based on the Nvidia Merlin dataloader (https://github.com/NVIDIA-Merlin/dataloader) which gives an easy-to-use API, uses the widely adopted Apache Parquet format to store data on disk and gives performant data loading with GPU-optimized data loaders that directly load the data from disk into GPU memory and then do a 0-copy transfer to PyTorch, TensorFlow or JAX (see Supp. Fig. 1 for details about data loading speed). The above-described data loading infrastructure was fast enough to fully utilize a Nvidia A100 GPU for the models trained in this paper. Moreover, Merlin comes with a wide range of supporting infrastructure like Docker containers (https://catalog.ngc.nvidia.com/orgs/nvidia/teams/merlin/containers/merlin-pytorch) from the NGC container hub which makes it easy for people to start using Merlin without the need to set up Python environments first.

### Data augmentation

The idea behind the data augmentation strategy developed in this paper is that the difference in raw gene space between the same cell type observed across two donors can be used as a data augmentation vector that can simulate how the gene expression of a cell might look like for a different donor. The general idea behind data augmentation is to have easy-to-compute transformations that can be applied during model training. Thus, in this case, we pre-compute augmentation vectors that can be added to the observed gene expression of a cell to artificially increase the training data size during model training:

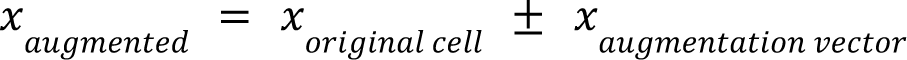

#### Calculation of augmentation vectors

The augmentation vectors are calculated as follows:

1. Subsample 500,000 cells from the training data to have an even distribution across cell types
2. Calculate the mean centroids grouped by cell type and donor
3. Calculate the difference vectors between the mean centroids from step 2 by cell type
4. Set all values in the range [-0.25, 0.25] to zero to enforce more sparse augmentation vectors
5. Clamp the resulting augmentation vectors to the interval of [-1.5, 1.5] to remove outlier values
6. Filter the resulting augmentation vectors for outliers by only sampling the used augmentation vectors from the most prominent k-means clusters (clustering is done with 50 clusters) → sample e.g. 5,000 augmentation vectors from the biggest k-means clusters (clusters with more than 2,000 difference vectors)

#### Calculation of augmented gene expression vectors during model training

The augmented gene expression vectors are calculated as follows:

1. Sample an augmentation vector *𝑥_augmentation vector_* from the set of augmentation vectors
2. Sample whether the augmentation vector is added to or subtracted from the original gene expression vector 𝑥_*original cell*_
3. Add/subtract the sampled augmentation vector to the original gene expression vector and clamp all values of the newly created vector to be within the interval of [0., 9.]

#### Explained variance by cell type before and after data augmentation

To estimate how our data augmentation influences the proportion of the overall variance that can be attributed to cell type variation, we fitted a linear regression (sci-kit learn LinearRegression) model which predicts the normalized gene expression based on the cell type and donor of each cell. This corresponds to the following design matrix:

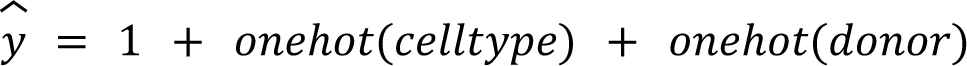

In the next step, the 𝑅^2^ score of the model fitted on the original/non-augmented data is compared to the one from the model fitted on the augmented data to show how the amount of total variation in gene expression, which can be attributed to the cell type, changes.

### Ontology-corrected cell type classification

The classification performance of the trained models in this paper is evaluated based on the macro average of the F1-scores for each individual cell type. The macro average is used to give each cell type the same weight in the overall classification performance. The F1-score is calculated as follows:

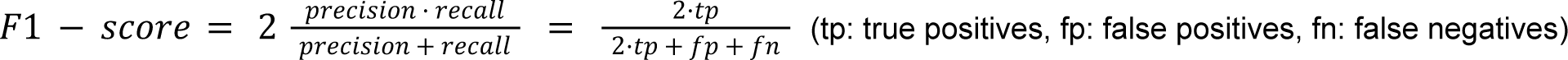

In order to deal with the often different granularity of annotations (e.g. label “T-cell” vs label “CD4-positive, alpha-beta T cell”) the following rules are applied to evaluate whether a prediction is considered right or wrong. A prediction is considered as right, either if the classifier predicts the same label as supplied by the original dataset, or if the classifier predicts a subtype of the label provided by the original dataset - we consider this as a right prediction as the prediction agrees with the true label up to the annotation granularity the author provided. The subtype relations are evaluated based on the Cell Ontology^29^. An example is if the model predicts the label “CD4-positive, alpha-beta T cell” when the author annotated cell type is “T cell”. Moreover, a prediction is considered wrong if the classifier predicts a parent cell type of the true label - we consider this as a wrong prediction as the author supplied a more fine-grained label that the classifier should replicate. An example is if the classifier predicts the label “T cell” while the cell is labeled as a “CD4-positive, alpha-beta T cell” in the original dataset. In all other cases, the prediction is considered wrong. Furthermore, the lookup of child nodes in the cell ontology is based on the Ontology Lookup Service (OLS): https://www.ebi.ac.uk/ols/ontologies/cl^29^

### Model details

#### scTab model

Our implementation of scTab is based on the TabNet architecture^31^ and is mostly taken from the dreamquark-ai/tabnet GitHub repository with some adaptation towards the single-cell use case. The input to the model is all 19,331 protein-coding genes (GENCODE v38/Ensembl 104) selected from the CELLxGENE census data. Moreover, unlike in the original TabNet model, we normalized the input data before feeding it into the neural network. scRNA-seq data is often normalized to have 10,000 counts per cell and is then log1p transformed afterward^7,12,23^, we applied the same normalization for our scTab model on top of the simple batch normalization layer, which is used in the original TabNet model to normalize the input features, as such a non-linear normalization cannot be achieved by a simple batch normalization layer.

The adapted TabNet architecture for scTab (Fig. 1b) consists of two key building blocks: The first building block is the feature transformer, which is a multi-layer perceptron with batch normalization (BN), skip connections, and a gated linear unit nonlinearity (GLU). On the one hand, the feature transformer is used to get a lower dimensional representation (described by dimensionality n_d) which is used to classify cell types. On the other hand, it is used to get a lower dimensional representation (described by dimensionality n_a) from which the feature attention mask is calculated. The feature attention mask is obtained by using a single linear layer followed by a batch normalization layer that maps from the attention embedding to the input feature space. The feature mask is then obtained by applying the 1.5-entmax^43^ function to the output of the linear projection layer. Using the 1.5-entmax function instead of the sparsemax function, which is used in the original TabNet model, improved training dynamics and yielded slightly higher model performance. The 1.5-entmax function is defined as follows:

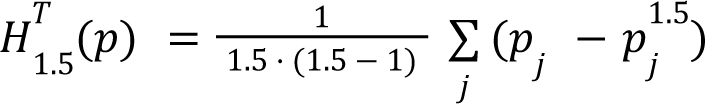

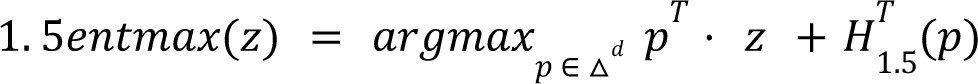

After obtaining the feature mask, the masked input features are fed into the feature transformer to obtain the feature embedding used to classify cell types. Thus, by giving the neural network the ability to mask individual input features, it can focus its network capacity only on more reliable input features. In contrast to the original TabNet model, we only used a single decision step as using more than one decision step only yielded marginal performance improvements and did not justify the increased computational costs.

The objective function used to train scTab is a cross-entropy loss where each cell type label is weighted in correspondence to its relative frequency in the training data to account for the strong class imbalance in the training data:

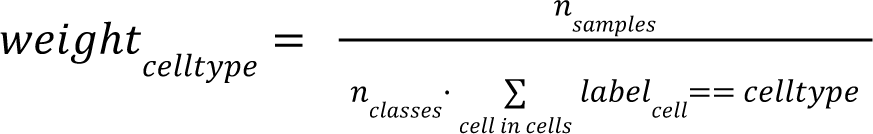

The model uncertainty is calculated based on deep ensembles ^32^ using 1 − 𝑚𝑎𝑥𝑖𝑚𝑢𝑚 𝑝𝑟𝑒𝑑𝑖𝑐𝑡𝑒𝑑 𝑝𝑟𝑜𝑎𝑏𝑖𝑙𝑖𝑡𝑦 as an estimate for the model uncertainty. Model uncertainties are calculated based on 5 ensemble models.

The models for Fig. 1 and Fig. 3 were fitted with our proposed data augmentation strategy. The models for Fig. 2 were fitted without data augmentation to better show the scaling with respect to the training data size.

List of used hyperparameters:

| Parameter | Value |
| --- | --- |
| batch_size | 2048 |
| learning_rate | 0.005 |
| learning rate scheduler | torch.optim.lr_scheduler.StepLR<br>gamma = 0.9<br>step_size = 1 epoch |
| optimizer | torch.optim.AdamW |
| weight_decay | 0.05 |
| n_d | 128 |
| n_a | 64 |
| n_shared | 3 |
| n_independent | 5 |
| n_steps | 1 |
| lambda_sparse | 1e-5 |
| mask_type | entmax |
| virtual_batch_size | 256 |
| augment_training_data | True |

#### XGBoost model

The input to the XGBoost model is a 256-dimensional PCA embedding due to the high memory usage and runtime of the XGBoost model. The PCA is only fitted on the training data to have a clear separation between the training and test set. Furthermore, the data is normalized to 10,000 counts per cell and is then log1p-transformed before calculating the PCA embeddings. The XGBoost model is fitted with the multi:softprob objective function and like for the scTab model classes are weighted in accordance to their relative frequency in the training data.

List of non-default hyperparameters:

| Parameter | Value |
| --- | --- |
| n_estimators | 800 |
| eta | 0.05 |
| subsample | 0.75 |
| max_depth | 10 |
| early_stopping_rounds | 10 |

For the benchmarks in this paper, we used XGBoost version 1.6.2

#### Multi-layer perceptron model (MLP)

The input to the model is all 19,331 protein-coding human genes selected from the CELLxGENE census data. The model is trained to predict the corresponding cell type label for each cell with a cross-entropy loss where each cell type is weighted in correspondence to its relative frequency (see scTab model).

The input count data is normalized to 10,000 counts per cell and is then log1p-transformed before feeding it into the model.

List of used hyperparameters:

| Parameter | Value |
| --- | --- |
| batch_size | 2048 |
| learning_rate | 0.002 |
| learning rate scheduler | torch.optim.lr_scheduler.StepLR<br>gamma = 0.9<br>step_size = 1 epoch |
| optimizer | torch.optim.AdamW |
| weight_decay | 0.05 |
| n_hidden | 8 |
| hidden_size | 128 |
| dropout | 0.1 |
| augment_training_data | True |

#### Linear model (ours)

The input to the model is all 19,331 protein-coding human genes selected from the CELLxGENE census data. The model consists of a single weight matrix and bias vector and is trained to predict the corresponding cell type label for each cell with a cross-entropy loss where each cell type is weighted in correspondence to its relative frequency (see scTab model).

The input count data is normalized to 10,000 counts per cell and is then log1p transformed before feeding them into the model.

List of used hyperparameters:

| Parameter | Value |
| --- | --- |
| batch_size | 2048 |
| learning_rate | 0.0005 |
| learning rate scheduler | torch.optim.lr_scheduler.StepLR<br>gamma = 0.9<br>step_size = 1 epoch |
| optimizer | torch.optim.AdamW |
| weight_decay | 0.01 |

#### CellTypist model

The CellTypist^7^ model was fitted in accordance with the best practice tutorial supplied on the CellTypist website with the difference that the mean centering step was disabled (*with_mean=False*) as this negatively impacted model performance and increased memory usage. Furthermore, the training data was subsampled to 1.5 million cells to keep both the memory usage (350GB of max memory) and runtime in check.

List of non-default hyperparameters:

| Parameter | Value |
| --- | --- |
| feature_selection | True |
| use_SGD | True |
| mini_batch | True |
| batch_number | 1500 |
| epochs | 10 |
| with_mean | False |

For the benchmarks in this paper, we used CellTypist version 1.5.3

## Code and data availability

GitHub - All code: https://github.com/theislab/scTab GitHub - Tutorials:

● Data loading tutorial: https://github.com/theislab/scTab/blob/main/notebooks-tutorials/data_loading.ipynb
● Loading trained models: https://github.com/theislab/scTab/blob/main/notebooks-tutorials/model_inference.ipynb

Data: https://pklab.med.harvard.edu/felix/data/merlin_cxg_2023_05_15_sf-log1p.tar.gz (164GB) Checkpoints: https://pklab.med.harvard.edu/felix/data/scTab-checkpoints.tar.gz (8.1GB)

## Acknowledgments

We thank the Cell Annotation Platform (CAP) team - especially Roman Mukhin, Andrey Isaev, and Uğur Bayındır - for their continuous support and feedback in developing and evaluating scTab, as well as Peter Kharchenko for his guidance on the implementation of new methods and computational frameworks on CAP. We thank Giovanni Palla, Luke Zappia, Alejandro Tejada Lapuerta, and Lukas Heumos for their suggestions that made the manuscript stronger. The authors also gratefully acknowledge the computational and data resources provided by the Leibniz Supercomputing Centre (www.lrz.de). The authors would also like to thank Keith Bayer and the Web and Advanced Research Platforms at Harvard Medical School for their support throughout this project.

D.S.F. acknowledges support from a German Research (DFG) fellowship through the Graduate School of Quantitative Biosciences Munich (QBM) [GSC 1006 to D.S.F.] and by the Joachim Herz Foundation. This work was supported in part by funding from the Eric and Wendy Schmidt Center at the Broad Institute of MIT and Harvard. A.C.V. acknowledges support from the National Institute of Health (DP2CA247831). This work was supported in part by funding from Schmidt Futures for the Cell Annotation Platform (to A.C.V. and F.J.T), as well as by the Helmholtz Association’s Initiative and Networking Fund on the HAICORE@FZJ partition.

This publication is part of the Human Cell Atlas – www.humancellatlas.org/publications

## Author contributions

F.F., D.S.F., and F.J.T. worked on pilot analyses and conceived the project. F.F. conducted the model implementation and analyses with input from E.B., D.S.F., and F.J.T. F.F. wrote and tested the software with E.B. and the Cell Annotation Platform development team. A.C.V. oversees and supports the Cell Annotation Platform effort. F.F., E.B., D.S.F., and F.J.T. wrote the manuscript. All authors discussed the results and commented on the manuscript.

## Supplements

**Supp. Table 1:** Classification performance of different models.

|  | F1-score (macro avg.) | Number of runs to calculate standard deviation |
| --- | --- | --- |
| scTab (deep learning) | 0.8295 ± 0.0007 | 5 |
| XGBoost (boosted decision trees) | 0.8127 ± 0.0005 | 5 |
| MLP (deep learning) | $0.7971 \pm 0.0012$ | 5 |
| Linear | $0.7848 \pm 0.0001$ | 4 |
| CellTypist (training data subsampled to 1.5 Mio cells) | $0.7304 \pm 0.0015$ | 4 |

**Supp. Table 2:** Performance of lung-specific versus cross-organ models evaluated on holdout test set subset to only lung-specific data.

|  | F1-score (macro avg.) on lung holdout data | Number of runs to calculate standard deviation |
| --- | --- | --- |
| scTab (cross-organ) | $0.7062 \pm 0.0122$ | 5 |
| scTab (lung only) | $0.7220 \pm 0.0078$ | 5 |
| Linear (cross-organ) | $0.5291 \pm 0.0041$ | 4 |
| Linear (lung only) | $0.7146 \pm 0.0040$ | 5 |

**Supp. Table 3:**
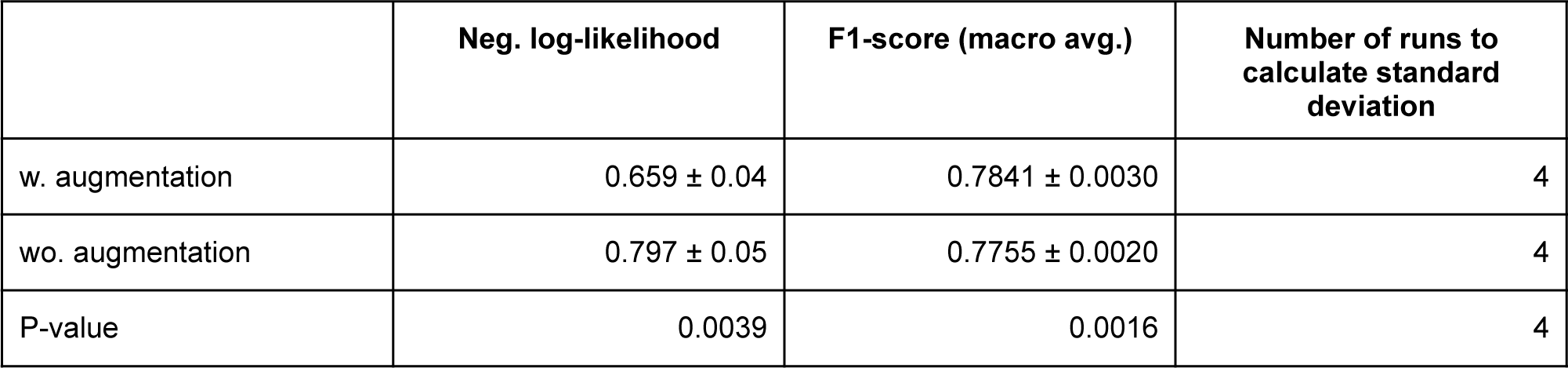
Effect of data augmentation on loss and F1-score (macro avg.) on holdout test set.

|  | Neg. log-likelihood | F1-score (macro avg.) | Number of runs to calculate standard deviation |
| --- | --- | --- | --- |
| w. augmentation | $0.659 \pm 0.04$ | $0.7841 \pm 0.0030$ | 4 |
| wo. augmentation | $0.797 \pm 0.05$ | $0.7755 \pm 0.0020$ | 4 |
| P-value | 0.0039 | 0.0016 | 4 |

**Supp. Table 4:** Classification performance of models with tuned versus default hyperparameters.

|  | F1-score (macro avg.) with default parameters | F1-score (macro avg.) with tuned parameters | Number of runs to calculate standard deviation |
| --- | --- | --- | --- |
| XGBoost | $0.5855 \pm 0.0112$ | $0.8127 \pm 0.0005$ | 4 |
| CellTypist | $0.6258 \pm 0.0036$ | $0.7304 \pm 0.0015$ | 4 |

**Supp. Table 5: Number of donors and cells per cell type and tissue combination.**

See *supp_table_donors_and_cells_per_tissue+cell_type.csv*

**Supp. Table 6: Number of shared tissues across individual cell types.**

See *supp_table_number_of_tisses_for_each_cell_type.csv*

**Supp. Table 7:** Total variation that can be attributed to the cell type before and after data augmentation.

| | Total variation attributed to cell type and donor ($R^2$) |
| --- | --- |
| original/non-augmented data | 0.189 |
| augmented data | 0.164 |

**Supp. Figure 1:**
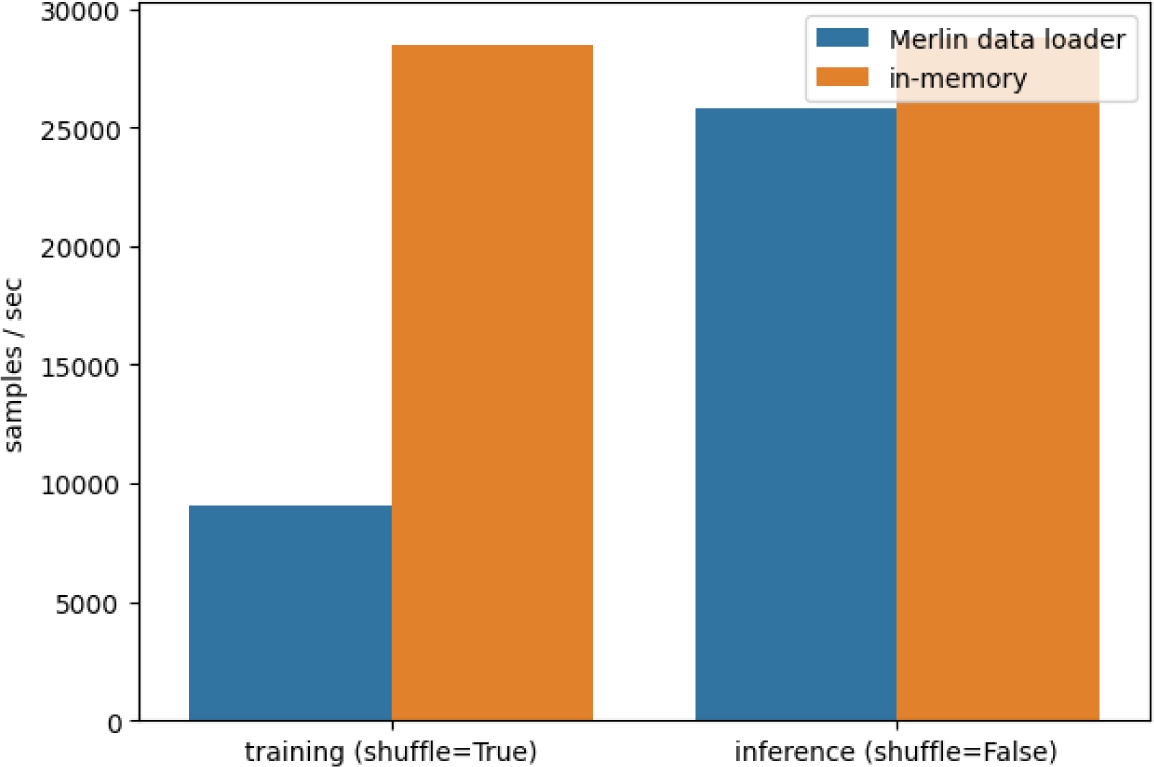
Data loading performance during model training (with data shuffling) and inference (without data shuffling). Benchmarks were run on a DGX-A100-320GB compute node with 14 cores and 80GB of memory allocated for the benchmark and half an A100 GPU (4g.20gb MIG). The training dataset consists of 15.2 million cells for the Merlin data loader and 1 million cells for the in-memory data loader. The validation dataset consists of 3.5 million cells for the Merlin data loader and 1 million cells for the in-memory data loader. Due to memory limitations for the in-memory data-loading, the training and validation set is subsampled to 1 million cells.

**Supp. Figure 2:**
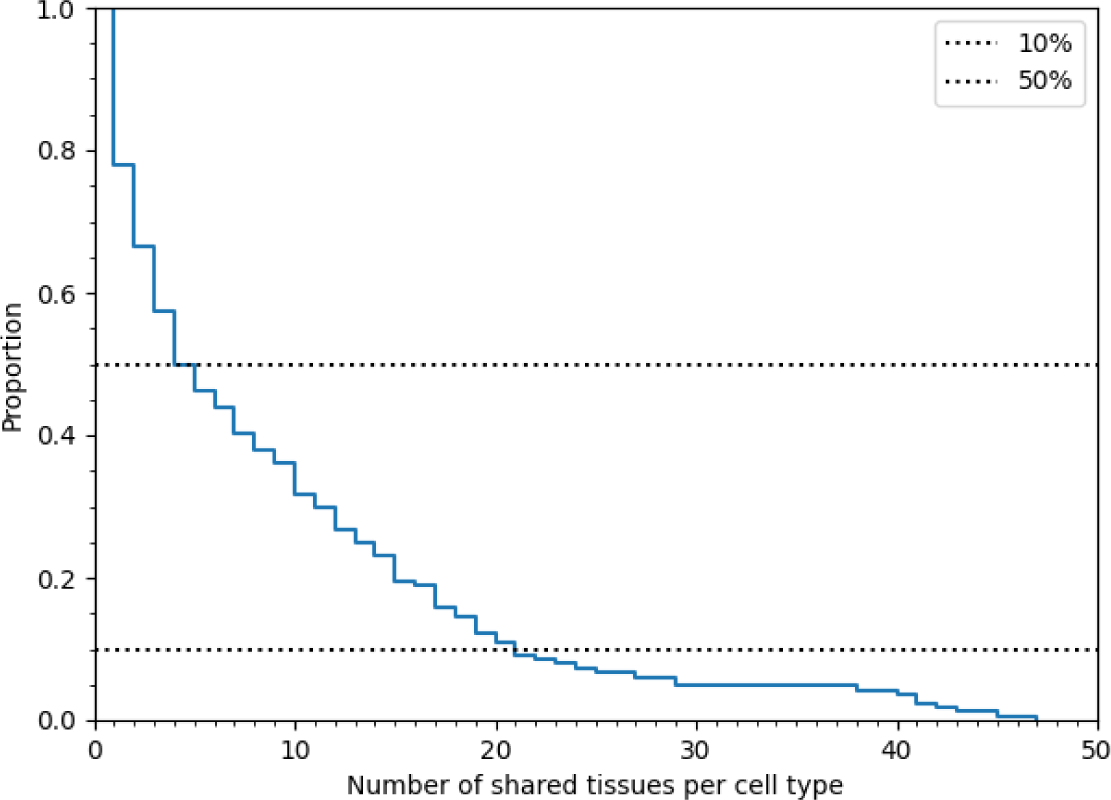
Number of shared tissues across individual cell types. The complementary empirical cumulative distribution function of how many tissues each cell type is observed over (see Supp Table 7 for a per cell type statistic).

**Supp. Figure 3:**
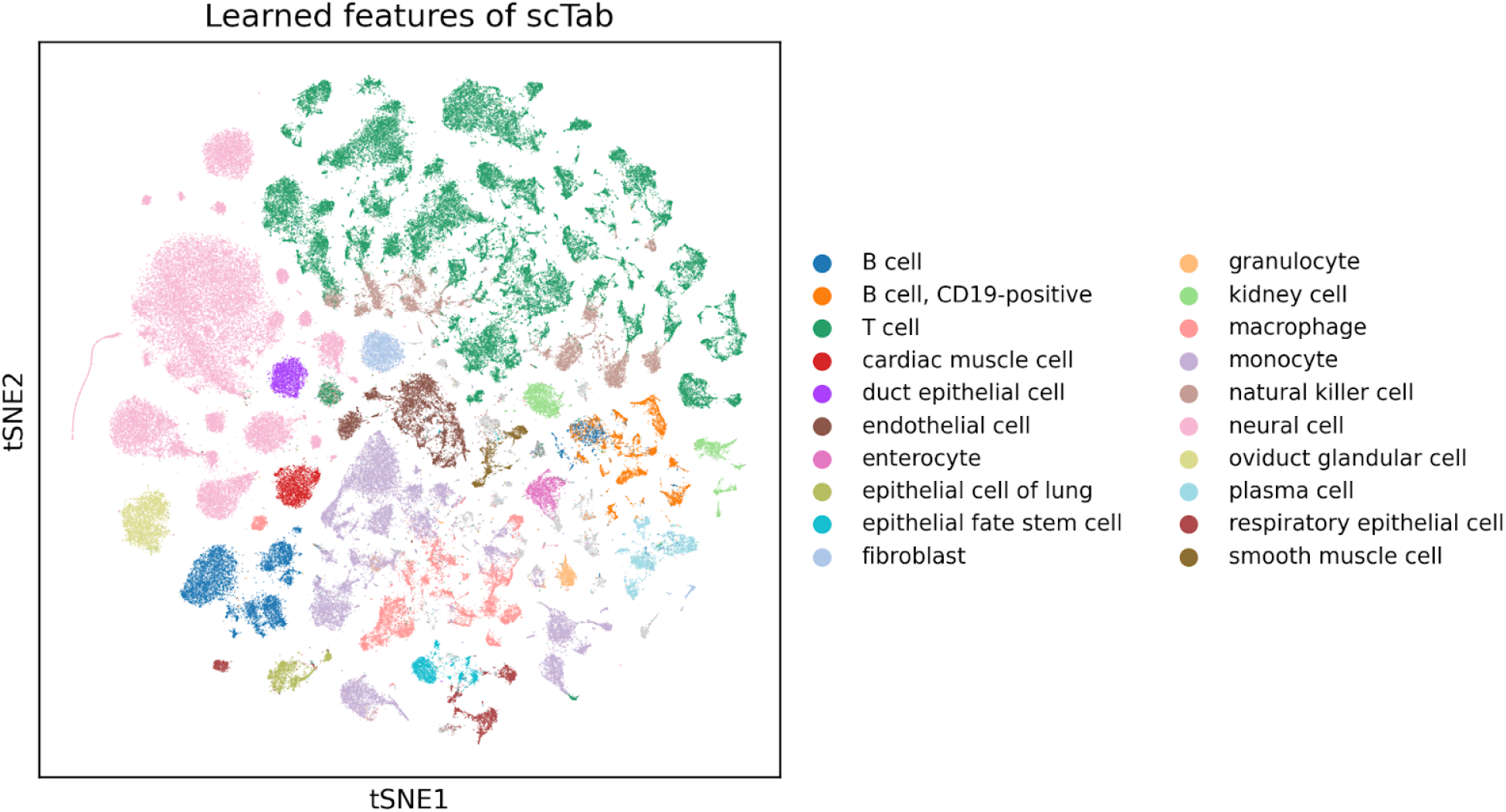
Learned features of scTab on holdout test data with granular cell type labels superimposed.

**Supp. Figure 4:**
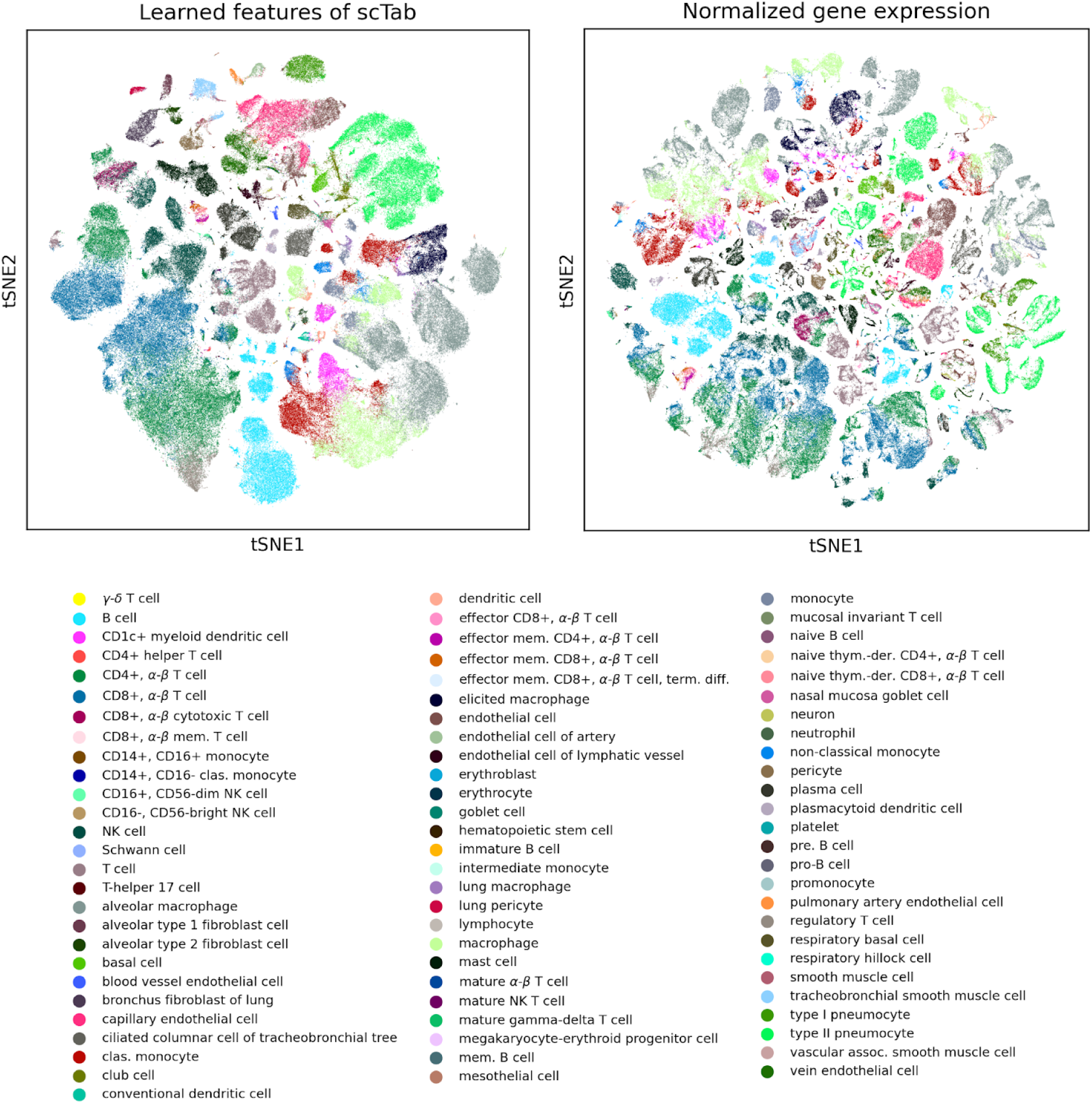
Learned features of scTab compared to the normalized gene expression of the input features on holdout test data subset to lung tissue only.

**Supp. Figure 5:**
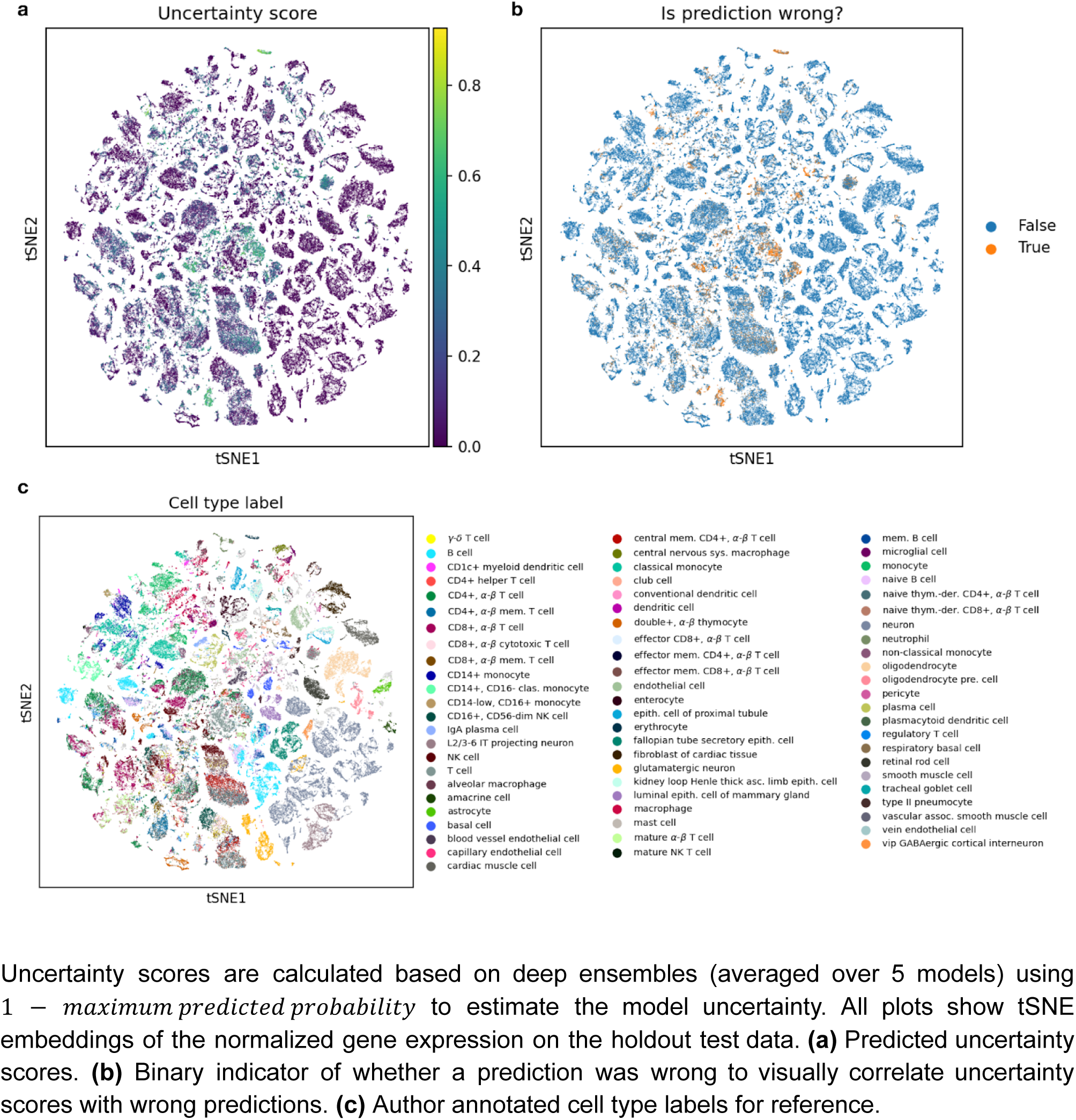
Uncertainty scores superimposed on tSNE plot of normalized gene expression on holdout test data. Uncertainty scores are calculated based on deep ensembles (averaged over 5 models) using 1 − 𝑚𝑎𝑥𝑖𝑚𝑢𝑚 𝑝𝑟𝑒𝑑𝑖𝑐𝑡𝑒𝑑 𝑝𝑟𝑜𝑏𝑎𝑏𝑖𝑙𝑖𝑡𝑦 to estimate the model uncertainty. All plots show tSNE embeddings of the normalized gene expression on the holdout test data. **(a)** Predicted uncertainty scores. **(b)** Binary indicator of whether a prediction was wrong to visually correlate uncertainty scores with wrong predictions. **(c)** Author annotated cell type labels for reference.

## References

1. Luecken, M. D. et al. Benchmarking atlas-level data integration in single-cell genomics. Nat. Methods 19, 41–50 (2022).

2. Luecken, M. D. & Theis, F. J. Current best practices in single-cell RNA-seq analysis: a tutorial. Mol. Syst. Biol. 15, e8746 (2019).

3. Heumos, L. et al. Best practices for single-cell analysis across modalities. Nat. Rev. Genet. 24, 550–572 (2023).

4. Amezquita, R. A. et al. Orchestrating single-cell analysis with Bioconductor. Nat. Methods 17, 137–145 (2020).

5. Abdelaal, T. et al. A comparison of automatic cell identification methods for single-cell RNA sequencing data. Genome Biol. 20, 194 (2019).

6. Köhler, N. D., Büttner, M., Andriamanga, N. & Theis, F. J. Deep learning does not outperform classical machine learning for cell-type annotation. bioRxiv (2019) doi:10.1101/653907.

7. Domínguez Conde, C., et al. Cross-tissue immune cell analysis reveals tissue-specific features in humans. Science 376, eabl5197 (2022).

8. Ergen, C., et al. Consensus prediction of cell type labels with popV. bioRxiv (2023) doi:10.1101/2023.08.18.553912.

9. Regev, A., et al. The Human Cell Atlas White Paper. arXiv [q-bio.TO] (2018).

10. Sikkema, L. et al. An integrated cell atlas of the lung in health and disease. Nat. Med. 29, 1563–1577 (2023).

11. Novella-Rausell, C., Grudniewska, M., Peters, D. J. M. & Mahfouz, A. A comprehensive mouse kidney atlas enables rare cell population characterization and robust marker discovery. bioRxiv 2022.07.02.498501 (2022) doi:10.1101/2022.07.02.498501.

12. Lopez, R., Regier, J., Cole, M. B., Jordan, M. I. & Yosef, N. Deep generative modeling for single-cell transcriptomics. Nat. Methods 15, 1053–1058 (2018).

13. Diehl, A. D. et al. The Cell Ontology 2016: enhanced content, modularization, and ontology interoperability. J. Biomed. Semantics 7, 44 (2016).

14. Fischer, D. S. et al. Sfaira accelerates data and model reuse in single cell genomics. Genome Biol. 22, 248 (2021).

15. Chan Zuckerberg CELLxGENE Discover. Cellxgene Data Portal https://cellxgene.cziscience.com/.

16. Clarke, Z. A. et al. Tutorial: guidelines for annotating single-cell transcriptomic maps using automated and manual methods. Nat. Protoc. 16, 2749–2764 (2021).

17. Lotfollahi, M. et al. Mapping single-cell data to reference atlases by transfer learning. Nat. Biotechnol. 40, 121–130 (2022).

18. Hao, Y. et al. Integrated analysis of multimodal single-cell data. Cell 184, 3573–3587.e29 (2021).

19. De Donno, C. et al. Population-level integration of single-cell datasets enables multi-scale analysis across samples. bioRxiv 2022.11.28.517803 (2022) doi:10.1101/2022.11.28.517803.

20. Huang, Y. & Zhang, P. Evaluation of machine learning approaches for cell-type identification from single-cell transcriptomics data. Brief. Bioinform. 22, (2021).

21. Cui, H., Wang, C., Maan, H. & Wang, B. scGPT: Towards Building a Foundation Model for Single-Cell Multi-omics Using Generative AI. *bioRxiv* 2023.04.30.538439 (2023) doi:10.1101/2023.04.30.538439.

22. Theodoris, C. V. et al. Transfer learning enables predictions in network biology. Nature 618, 616–624 (2023).

23. Heimberg, G. et al. Scalable querying of human cell atlases via a foundational model reveals commonalities across fibrosis-associated macrophages. bioRxiv 2023.07.18.549537 (2023) doi:10.1101/2023.07.18.549537.

24. Xu, C. et al. Probabilistic harmonization and annotation of single-cell transcriptomics data with deep generative models. Mol. Syst. Biol. 17, e9620 (2021).

25. Shwartz-Ziv, R. & Armon, A. Tabular data: Deep learning is not all you need. (2021) doi:10.48550/ARXIV.2106.03253.

26. Kaplan, J., et al. Scaling Laws for Neural Language Models. arXiv [cs.LG] (2020) doi:10.48550/ARXIV.2001.08361.

27. Krizhevsky, A., Sutskever, I. & Hinton, G. E. Imagenet classification with deep convolutional neural networks. Adv. Neural Inf. Process. Syst. 25, (2012).

28. Shorten, C. & Khoshgoftaar, T. M. A survey on image data augmentation for deep learning. J. Big Data 6, (2019).

29. Jupp, S., Burdett, T., Leroy, C. & Parkinson, H. E. A new Ontology Lookup Service at EMBL-EBI. SWAT4LS 2, 118–119 (2015).

30. Osumi-Sutherland, D. et al. Cell type ontologies of the Human Cell Atlas. Nat. Cell Biol. 23, 1129–1135 (2021).

31. Arik, S. O. & Pfister, T. TabNet: Attentive Interpretable Tabular Learning. (2019) doi:10.48550/ARXIV.1908.07442.

32. Lakshminarayanan, B., Pritzel, A. & Blundell, C. Simple and Scalable Predictive Uncertainty Estimation using Deep Ensembles. arXiv [stat.ML] (2016).

33. Zhang, C., Bengio, S., Hardt, M., Recht, B. & Vinyals, O. Understanding deep learning (still) requires rethinking generalization. Commun. ACM 64, 107–115 (2021).

34. Deng, J. et al. ImageNet: A large-scale hierarchical image database. in 2009 IEEE Conference on Computer Vision and Pattern Recognition (IEEE, 2009). doi:10.1109/cvpr.2009.5206848.

35. Russakovsky, O. et al. ImageNet Large Scale Visual Recognition Challenge. arXiv [cs.CV] (2014).

36. Lin, T.-Y. et al. Microsoft COCO: Common Objects in Context. in Computer Vision – ECCV 2014 740–755 (Springer International Publishing, 2014).

37. Wang, A., et al. GLUE: A Multi-Task Benchmark and Analysis Platform for Natural Language Understanding. arXiv [cs.CL] (2018).

38. Wang, A., et al. SuperGLUE: A Stickier Benchmark for General-Purpose Language Understanding Systems. arXiv [cs.CL] (2019).

39. Luong, M.-T. & Manning, C. Stanford neural machine translation systems for spoken language domains. in Proceedings of the 12th International Workshop on Spoken Language Translation: Evaluation Campaign 76–79 (aclanthology.org, dec # ‘ 3-4’, year = ‘2015’, address = ‘Da Nang, Vietnam’, url = ‘https:/aclanthology.org/2015.iwslt-evaluation.11’, pages = ‘76--79’ 2015).

40. Hao, M., et al. Large scale foundation model on single-cell transcriptomics. bioRxiv (2023) doi:10.1101/2023.05.29.542705.

41. Yang, F. et al. scBERT as a large-scale pretrained deep language model for cell type annotation of single-cell RNA-seq data. Nature Machine Intelligence 4, 852–866 (2022).

42. Grill, J.-B., et al. Bootstrap your own latent: A new approach to self-supervised Learning. *arXiv [cs.LG]* (2020) doi:10.48550/ARXIV.2006.07733.

43. Peters, B., Niculae, V. & Martins, A. F. T. Sparse sequence-to-sequence models. in Proceedings of the 57th Annual Meeting of the Association for Computational Linguistics (Association for Computational Linguistics, 2019). doi:10.18653/v1/p19-1146.

